# Imbalanced Nucleocytoskeletal Connections Create Common Polarity Defects in Progeria and Physiological Aging

**DOI:** 10.1101/497297

**Authors:** Wakam Chang, Yuexia Wang, G.W. Gant Luxton, Cecilia Östlund, Howard J. Worman, Gregg G. Gundersen

## Abstract

Studies of the accelerated aging disorder Hutchinson-Gilford progeria syndrome (HGPS) can potentially reveal cellular defects associated with physiological aging. HGPS results from expression and abnormal nuclear envelope association of a farnesylated, truncated variant of prelamin A called progerin. We surveyed the diffusional mobilities of nuclear membrane proteins to identify proximal effects of progerin expression. The mobilities of three proteins were reduced in fibroblasts from children with HGPS compared to normal fibroblasts: SUN2, nesprin-2G, and emerin. These proteins function together in nuclear movement and centrosome orientation in fibroblasts polarizing for migration. Both processes were impaired in fibroblasts from children with HGPS and in NIH3T3 fibroblasts expressing progerin, but were restored by inhibiting protein farnesylation. Progerin affected both the coupling of the nucleus to actin cables and the oriented flow of the cables necessary for nuclear movement and centrosome orientation. Progerin overexpression increased levels of SUN1, which couples the nucleus to microtubules through nesprin-2G and dynein, and microtubule association with the nucleus. Reducing microtubule-nuclear connections through SUN1 depletion or dynein inhibition rescued the polarity defects. Nuclear movement and centrosome orientation were also defective in fibroblasts from normal individuals over 60 years old and both defects were rescued by reducing the increased level of SUN1 in these cells or inhibiting dynein. Our results identify imbalanced nuclear engagement of the cytoskeleton (microtubules, high; actin filaments, low) as the basis for intrinsic cell polarity defects in HGPS and physiological aging and suggest that rebalancing the connections can ameliorate the defects.

**Significance:** The rare, premature aging syndrome HGPS arises from expression of a pathological prelamin A variant, termed progerin. Studies of progerin may identify treatments for HGPS and reveal novel cellular and molecular characteristics of normal aging. Here, we show that progerin selectively affects mobilities of three nuclear membrane proteins, SUN2, nesprin-2G and emerin that position the nucleus and establish cell polarity essential for migration. We find that both processes are defective in fibroblasts from children with HGPS and aged (> 60 years) individuals. The mechanism underlying these defects is excessive interaction of the nucleus with microtubules. Our work identifies nuclear-based defects in cell polarization as intrinsic factors in premature and physiological aging and suggests means for correcting them.

## Introduction

Hutchinson-Gilford progeria syndrome (HGPS) is an accelerated aging syndrome caused by mutations in the *LMNA* gene encoding prelamin A and lamin C (1, 2). In normal cells, prelamin A undergoes a series of modifications to produce mature lamin A. The C-terminal CaaX motif of prelamin A is farnesylated, followed by C-terminal methylation and removal of the last three amino acids, and a final cleavage that removes another 15 amino acids of the C-terminus including the farnesylated cysteine (3). In HGPS, a cryptic splice site in prelamin A mRNA is activated resulting in the production of a truncated variant, termed progerin, which lacks the final cleavage site and remains farnesylated (1, 2). By retaining its farnesyl moiety, progerin accumulates on the inner nuclear membrane where it affects nuclear architecture and functions associated with the nuclear lamina (4–6). Progerin expression causes nuclear shape abnormalities and alters many nuclear functions and cellular pathways (4, 7–9). In most cases, how progerin expression leads to these alterations is poorly understood.

Most studies have attributed alterations caused by progerin to its effects on the lamina. Yet, because progerin associates with the inner nuclear membrane, it may also dominantly interfere with other nuclear envelope proteins (10). Lamin A plays a critical role in the function of the linker of nucleoskeleton and cytoskeleton (LINC) complex. This complex is composed of inner nuclear membrane SUN proteins and outer nuclear membrane KASH proteins (known as nesprins in vertebrates) (11, 12). Through nesprins’ interaction with the cytoskeleton, the LINC complex contributes to nuclear movement and positioning, organization of the cytoskeleton, mechanotransduction to the nucleus, DNA repair and meiotic chromosome movements (11–16). Lamin A interacts with both SUN1 and SUN2, the major SUN domain proteins in somatic cells. Although it is not critical for their nuclear localization, it affects their mobilities and lack of lamin A prevents the anchoring of the LINC complex that is necessary for transmitting force (17–19).

There are few studies of the effects of progerin on the cellular functions of LINC complex. Progerin, like farnesylated prelamin A, exhibits increased association with SUN1 compared to SUN2 (17, 20). This may explain the increased levels of SUN1 observed in fibroblasts from children with HGPS and is likely to have deleterious physiological consequences as skeletal phenotypes and shortened life span of progeroid mouse models are improved by knocking out SUN1 (21). In homeostatic positioning of nuclei, SUN1 and SUN2 function separately to support nesprin-2G coupling to microtubules and actin filaments, respectively, and overexpressing one of the SUN proteins interferes with the function of the other (“transdominant inhibition”) (22). Thus, the upregulation of SUN1 in fibroblasts from individuals with HGPS may itself alter LINC complex function.

Here, we explore the hypothesis that progerin expression alters nuclear membrane proteins through its association with the inner nuclear membrane. We identify a subset of nuclear membrane proteins that are altered by progerin expression and show that their function in nuclear movement and cell polarity is disrupted. We find similar defects in fibroblasts from aged individuals and identify excessive microtubule interactions with the nucleus as the cause in fibroblasts from both HGPS and aged individuals.

## Results

### Progerin expression reduces the diffusional mobilities of selected nuclear membrane proteins

We surveyed the diffusional mobilities of EGFP-tagged integral nuclear membrane proteins by fluorescence recovery after photobleaching (FRAP) in fibroblasts from children with HGPS (HGPS fibroblasts) and age and sex matched controls (SI Appendix, Table S1). We tested proteins that interact directly or indirectly with lamins A and C (lamin A/C), including emerin, lamina-associated polypeptide 1 (LAP1), and LINC complex proteins (12, 23, 24). This survey revealed that the mobilities of mini-nesprin-2G (mN2G), a truncated form of nesprin-2G that functionally supports actin-dependent nuclear movement (25), SUN2, and emerin, but not five other nuclear membrane proteins (nesprin-3α, nesprin-3β, nesprin-4, SUN1 or LAP1) were significantly reduced in fibroblasts from individuals with HGPS compared to those from unaffected controls (Fig. 1A; SI Appendix, Fig. S1A,B). In HGPS fibroblasts from four different individuals, the halftimes of recovery (t_1/2_) of mN2G and SUN2 were consistently longer than those in control fibroblasts (SI Appendix, Fig. S1C). We did not detect a reduction in the mobility of SUN1, previously reported in HeLa cells overexpressing progerin (20), in all four HGPS fibroblast samples we tested (SI Appendix, Fig. S1C). Despite their altered diffusional mobilities, the nuclear localizations of nesprin-2G, SUN2, and emerin appeared similar in HGPS and control fibroblasts (SI Appendix, Fig. S2A).

**Figure 1.**
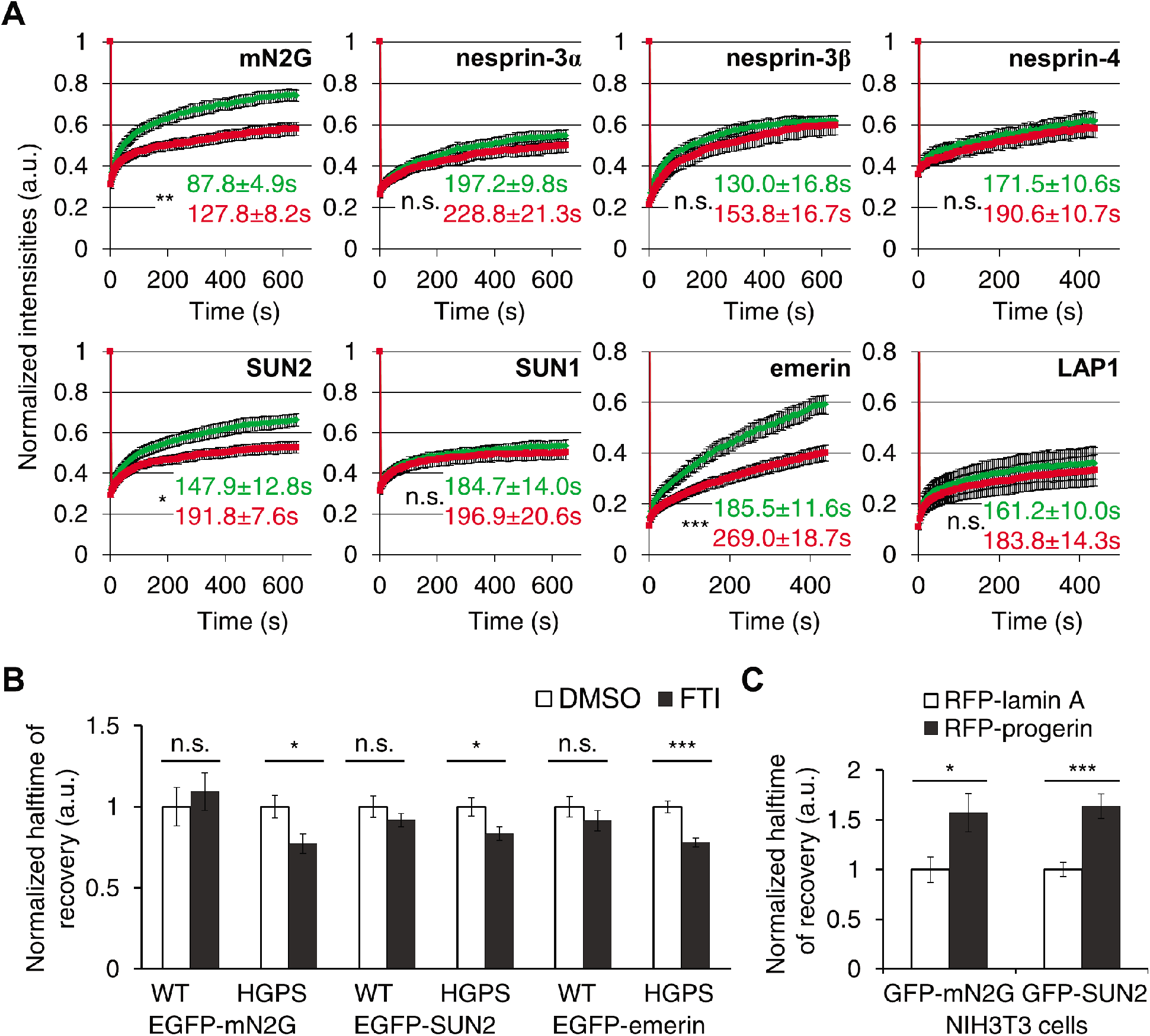
Progerin expression reduces the diffusional mobilities of a subset of nuclear envelope proteins. *(A)* Normalized FRAP in the nuclear envelope for indicated EGFP-tagged proteins in control (WT, green) and HGPS fibroblasts (red). Numbers are t_1/2_ of FRAP calculated and used for statistical tests. *(B)* Normalized t_1/2_ of FRAP of EGFP-tagged mN2G, SUN2, and emerin in HGPS fibroblasts treated with 2.5 μM FTI-277. Values are normalized to cells treated with dimethyl sulfoxide (DMSO). *(C)* Normalized t_1/2_ of FRAP of EGFP-tagged mN2G and SUN2 in NIH3T3 fibroblasts expressing RFP-tagged lamin A or progerin. See SI Appendix, Table S2, S3 for absolute numbers. Data are mean ± SEM from ≥ 3 experiments (*n* > 15 cells). n.s. *p* > 0.05, \**p* < 0.05, \*\**p* < 0.01, \*\*\**p* < 0.001 by Student’s t-test.

We next tested whether progerin was responsible for the altered mobilities. Treatment of HGPS fibroblasts with a protein farnesyltransferase inhibitor (FTI-277) at a concentration that blocked prelamin A processing (SI Appendix, Fig. S2B) significantly increased the diffusional mobilities and decreased the t_1/2_ of EGFP-tagged mN2G, SUN2, and emerin in HGPS fibroblasts without affecting their baseline mobilities in normal fibroblasts (Fig. 1B, SI Appendix, S1D). SUN protein levels were not affected by FTI treatment (SI Appendix, Fig. S2C). Ectopic expression of RFP-progerin in NIH3T3 fibroblasts also reduced the diffusional mobilities of mN2G and SUN2 (Fig. 1C, SI Appendix, S1E). Thus, progerin farnesylation was necessary and progerin expression was sufficient to reduce the diffusional mobilities of mN2G and SUN2.

### Progerin expression causes polarity defects in migratory fibroblasts

Nesprin-2G, SUN2, and emerin are all required for rearward nuclear movement and centrosome positioning in fibroblasts and myoblasts polarizing for migration (25–27), which is known to be impaired by HGPS (7, 28). After stimulation with serum or the serum factor lysophosphatidic acid (LPA), nesprin-2G and SUN2 assemble into transmembrane actin-associated nuclear (TAN) lines that couple the nucleus to perinuclear actin cables to move it to the cell rear resulting in centrosome orientation (25, 29). In contrast, emerin interacts with myosin-IIB on the outer nuclear membrane to orient the flow of perinuclear actin cables (26). In LPA-stimulated HGPS fibroblasts, rearward nuclear positioning was impaired, preventing proper centrosome orientation; however, these processes were restored by treatment with FTI-277 (Fig. 2A-C). To test if progerin was directly responsible for these defects, we expressed EGFP-tagged lamin A variants in NIH3T3 fibroblasts. Expression of EGFP-tagged progerin, but not wild type lamin A or a nonfarnesylatable progerin variant whose carboxyl-terminal CSIM motif was mutated to SSIM (30), inhibited rearward nuclear positioning and centrosome orientation (Fig. 2D-F). Treatment with FTI-277 reversed the effects of EGFP-progerin but had no effects in cells expressing EGFP-lamin A (Fig. 2D-F). Thus, farnesylated progerin inhibits the acquisition of polarity in fibroblasts by preventing rearward nuclear positioning.

**Figure 2.**
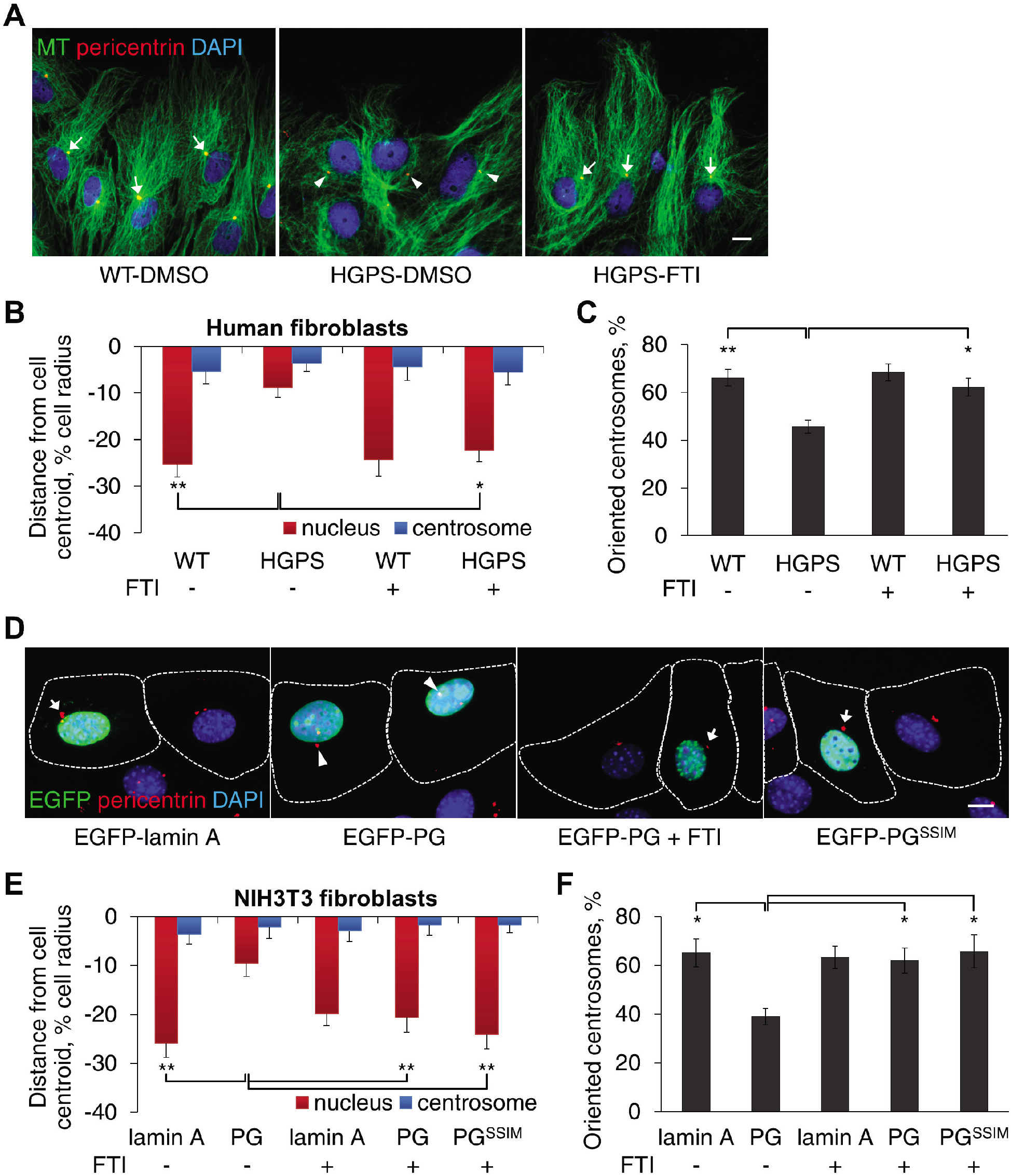
Progerin expression causes polarity defects in migratory fibroblasts. *(A)* Representative images of microtubules (MT), centrosomes (pericentrin), and nuclei (DAPI; 4’,6-diamidino-2-phenylindole) in LPA-stimulated control (WT) and HGPS fibroblasts pretreated with vehicle (DMSO) or FTI-277. Arrows, oriented centrosomes; arrowheads, non-oriented centrosomes. *(B)* Quantification of nuclear (red) and centrosomal (blue) positions along the front-back axis of LPA-stimulated WT or HGPS fibroblasts treated with or without FTI-277. The cell centroid is defined as “0”; + values, toward the leading edge; – values, toward the cell rear. *(C)* Quantification of centrosome orientation for the cells treated as in B. Random centrosome orientation is 33%. *(D)* Representative images of LPA-stimulated NIH3T3 fibroblasts transiently expressing EGFP-tagged lamin A, progerin (PG, +/− FTI treatment) or a non-farnesylatable progerin variant (PG^SSIM^). Dotted lines, cell outlines; arrows and arrowheads as in A. *(E,F)* Quantification of nuclear and centrosomal positions *(E)* and centrosome orientation *(F)* in LPA-stimulated NIH3T3 fibroblasts cells expressing the indicated EGFP-tagged constructs and treated with or without FTI-277. Data are mean ± SEM from ≥ 3 experiments (*n* > 90 cells per experiment). \**p* < 0.05, \*\**p* < 0.01 by t-test. Bars, A,C: 10 μm.

### Progerin expression affects TAN line anchorage and actin retrograde flow

To determine how progerin affects nuclear positioning in HGPS fibroblasts, we examined TAN line assembly and actin retrograde flow. TAN lines, which are identified by the accumulation of EGFP-mN2G and endogenous SUN2 along perinuclear actin cables on the dorsal nuclear surface (25), formed similarly in HGPS and control fibroblasts (Fig. 3A-C). However, live cell imaging revealed that TAN lines in cells expressing EGFP-progerin moved rearward, albeit slowly (see rate of actin retrograde flow below), on immobile nuclei (Fig. 3C, SI Appendix, Video S1). This slippage of TAN lines indicated that they are not strongly anchored in these cells, a phenotype similar to that occurring in fibroblasts lacking lamin A/C (19).

**Figure 3.**
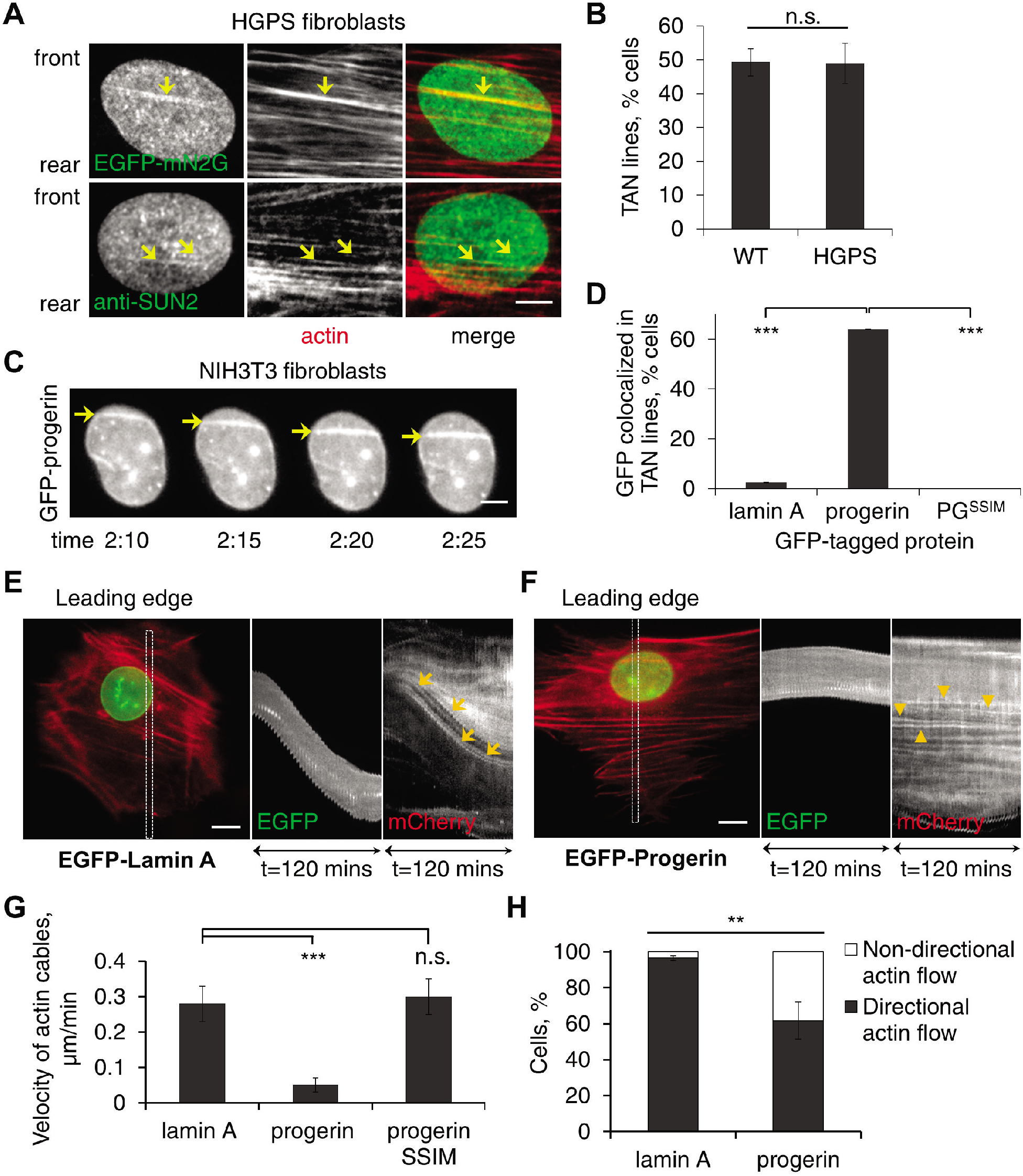
Progerin expression affects TAN line anchorage to the nuclear envelope and retrograde actin flow. *(A)* Representative images of a nucleus in an LPA-stimulated HGPS fibroblast expressing EGFP-mN2G (top) or stained for SUN2 (bottom). mN2G and SUN2 formed linear structures (arrows) that colocalized with actin filaments (stained by phalloidin) indicating TAN line formation. *(B)* Quantification of TAN line formation in control (WT) and HGPS fibroblasts (N = 3 experiments, *n* > 70 cells). *(C)* Images from a movie of EGFP-progerin in NIH3T3 fibroblasts. Note accumulation of EGFP-progerin in TAN lines (arrows) that slip over an immobile nucleus. (See SI Appendix, Video S1 for mCherry-LifeAct signal). Time, h:min after LPA treatment. *(D)* Quantification of cells with GFP-positive TAN lines (N = 3 experiments, *n* > 50 cells). *(E,F)* Representative kymographs from movies of LPA-stimulated NIH3T3 fibroblasts expressing EGFP-lamin A (E, from SI Appendix, Video S2) or EGFP-progerin (F, from SI Appendix, Video S3) together with mCherry-Lifeact. The color image is the first frame of the movies (30 min after LPA stimulation) and shows the region (box) used for the kymographs on the right. Note slower rearward actin flow in cells expressing EGFP-progerin (arrowheads) compared to EGFP-lamin A (arrows). Leading edge of the cells is toward the top in all panels. *(G)* Quantification of the velocity of retrograde actin flow near the leading edge in NIH3T3 fibroblasts expressing mCherry-LifeAct and EGFP-tagged proteins as indicated (N = 3 experiments, *n* > 40 actin cables). *(H)* Quantification of the directionality of actin flow in NIH3T3 fibroblasts expressing mCherry-LifeAct and indicated EGFP-tagged proteins (N = 3 experiments, *n* > 50 cells). Data are mean ± SEM from ≥ 3 experiments. n.s. *p* > 0.05, \*\**p* < 0.01, \*\*\**p* < 0.001 by t-test. Bars, A,C: 5 μm, E,F: 10 μm.

Expression of EGFP-progerin also affected the retrograde flow of actin near the leading edge. As previously reported (19, 25), in control NIH3T3 fibroblasts and those expressing EGFP-lamin A, actin cables near the leading edge exhibited predominantly rearward movement at a rate of 0.28 ± 0.05 μm/min (Fig. 3E and SI Appendix, Video S2). In contrast, in cells expressing EGFP-progerin the velocity was only 0.05 ± 0.02 μm/min (Fig. 3F and SI Appendix, Video S3). The velocity of actin cables in cells expressing a nonfarnesylatable progerin variant was 0.30 ± 0.05 μm/min (Fig. 3G). Actin cables in cells expressing EGFP-progerin also exhibited significantly more non-directional actin flow compared to cells expressing EGFP-lamin A (Fig. 3H). Thus, farnesylated progerin affects rearward nuclear movement by both weakening TAN line anchorage and disturbing retrograde actin flow.

### Cell polarity defects in HGPS are mediated by increased SUN1

The defects in TAN line anchorage and actin retrograde flow in progerin overexpressing cells are reminiscent of those in lamin A/C deficient cells (19) and emerin deficient cells (26), respectively. How does progerin affect both these processes? We considered the possibility that the upregulation of SUN1 observed with progerin overexpression (21) may exert a transdominant inhibitory effect preventing SUN2-nesprin-2G coupling to the actin cytoskeleton (22). We confirmed the previously reported upregulation of SUN1 in HGPS fibroblasts (21) and additionally found that stable expression of progerin, but not lamin A, in NIH3T3 fibroblasts proportionately increased the level of SUN1 but not SUN2 in the nucleus (Fig. 4A, B). This shows that progerin overexpression specifically increases SUN1 levels and is consistent with earlier reports that the levels of SUN1 and SUN2 are not interdependent (22, 25). These findings suggest that an elevated level of SUN1 may contribute to the nuclear movement and cell polarity defects in HGPS fibroblasts. Indeed, stable overexpression of SUN1 in NIH3T3 fibroblasts prevented rearward nuclear movement and centrosome orientation (Fig. 4C, D). Critically, SUN1 depletion by siRNA treatment (SI Appendix, Fig. S3A) rescued the defects in rearward nuclear movement and centrosome orientation in HGPS fibroblasts (Fig. 4E-G). As with FTI-277 treatment, SUN1 depletion from HGPS fibroblasts significantly increased the diffusional mobility of EGFP-mN2G (SI Appendix, Fig. S3B, C). These results indicate that elevated SUN1 expression is responsible for the cell polarity defects in HGPS fibroblasts.

**Figure 4.**
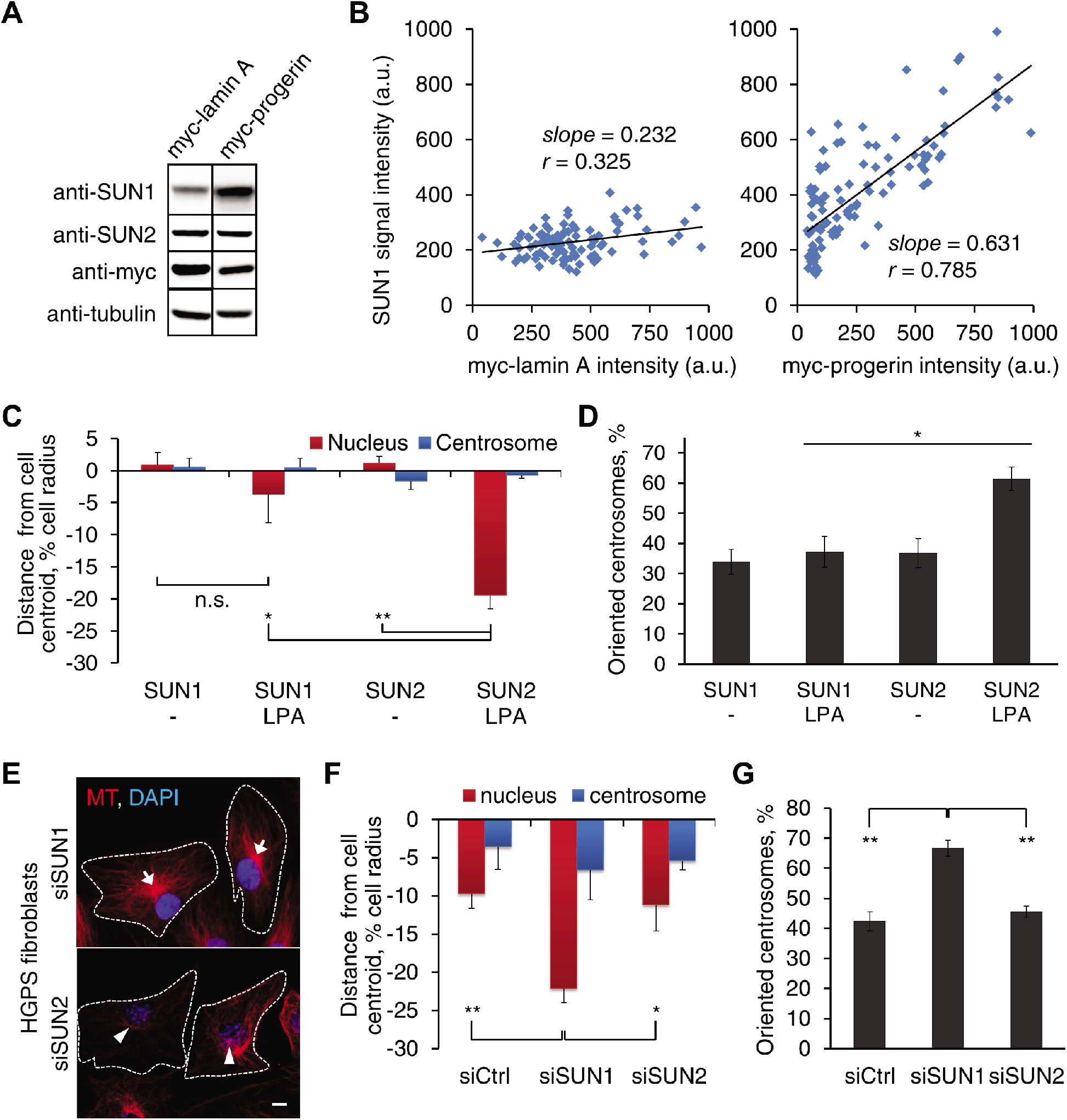
Cell polarity defects in HGPS fibroblasts are mediated by increased SUN1. *(A)* Immunoblot of lysates from myc-lamin A or progerin expressing NIH3T3 fibroblasts probed with the indicated antibodies. Tubulin is a loading control. *(B)* Quantification of signal intensities in individual NIH3T3 fibroblasts (n > 120) stably expressing myc-tagged lamin A (left) or progerin (right) and stained for myc and SUN1. Correlation coefficients (r) and slopes were calculated assuming a linear relationship. *(C,D)* Quantification of nuclear and centrosomal positions *(C)* and centrosome orientation *(D)* in NIH3T3 fibroblasts stably expressing myc-SUN1 or myc-SUN2 and treated with or without LPA. *(E)* Representative images of microtubules (MT) and nuclei (DAPI) in LPA-stimulated HGPS fibroblasts treated with indicated siRNAs. Arrows, oriented centrosomes; arrowheads, non-oriented centrosomes; dotted lines, cell borders. Bar, 10 μm. *(F,G)* Quantification of nuclear and centrosomal positions *(F)* and centrosome orientation *(G)* in LPA-stimulated HGPS fibroblasts treated with indicated siRNAs. Data are mean ± SEM from *n* ≥ 3 experiments *(n* > 90 cells). n.s. *p* > 0.05, \**p* < 0.05, \*\**p* < 0.01 by t-test.

### Cell polarity defects in HGPS fibroblasts are mediated by the microtubule cytoskeleton

SUN1 and SUN2 are ubiquitously expressed in somatic cells but separately support nesprin-2G dependent nuclear movement by microtubules and actin filaments, respectively (22). Additionally, overexpression of one SUN protein transdominantly inhibits the ability of the other to support nuclear movement (Fig. 4C and ref. 22). The elevated levels of SUN1 and the rescue of actin- and SUN2-dependent rearward nuclear positioning by SUN1 depletion, suggested that the nucleus in HGPS fibroblasts may interact abnormally with microtubules. Indeed, we observed excessive association of microtubules with nuclei in HGPS fibroblasts compared to control fibroblasts (Fig. 5A). To directly test if microtubules contribute to polarity defects in HGPS fibroblasts, we used a dynein inhibitor, HPI-4 (ciliobrevin A), which inhibits SUN1- and microtubule-dependent nuclear movement in fibroblasts (22). At a low concentration, HPI-4 did not improve nuclear morphology but rescued actin-dependent nuclear movement in both HGPS fibroblasts (Fig. 5B, C) and NIH3T3 fibroblasts overexpressing progerin (Fig. 5E, F). HPI-4 also increased centrosome orientation (Fig. 5D, G), even though it caused the centrosome to move rearward with the nucleus as expected given dynein’s role in maintaining the centrosome in the cell center (29). HPI-4 treatment did not affect SUN levels and did not rescue the mobilities of mN2G and SUN2 (SI Appendix, Fig. S3D-F), suggesting that the bulk mobility of these proteins is determined by their nuclear anchorage rather than their cytoskeletal interactions.

**Figure 5.**
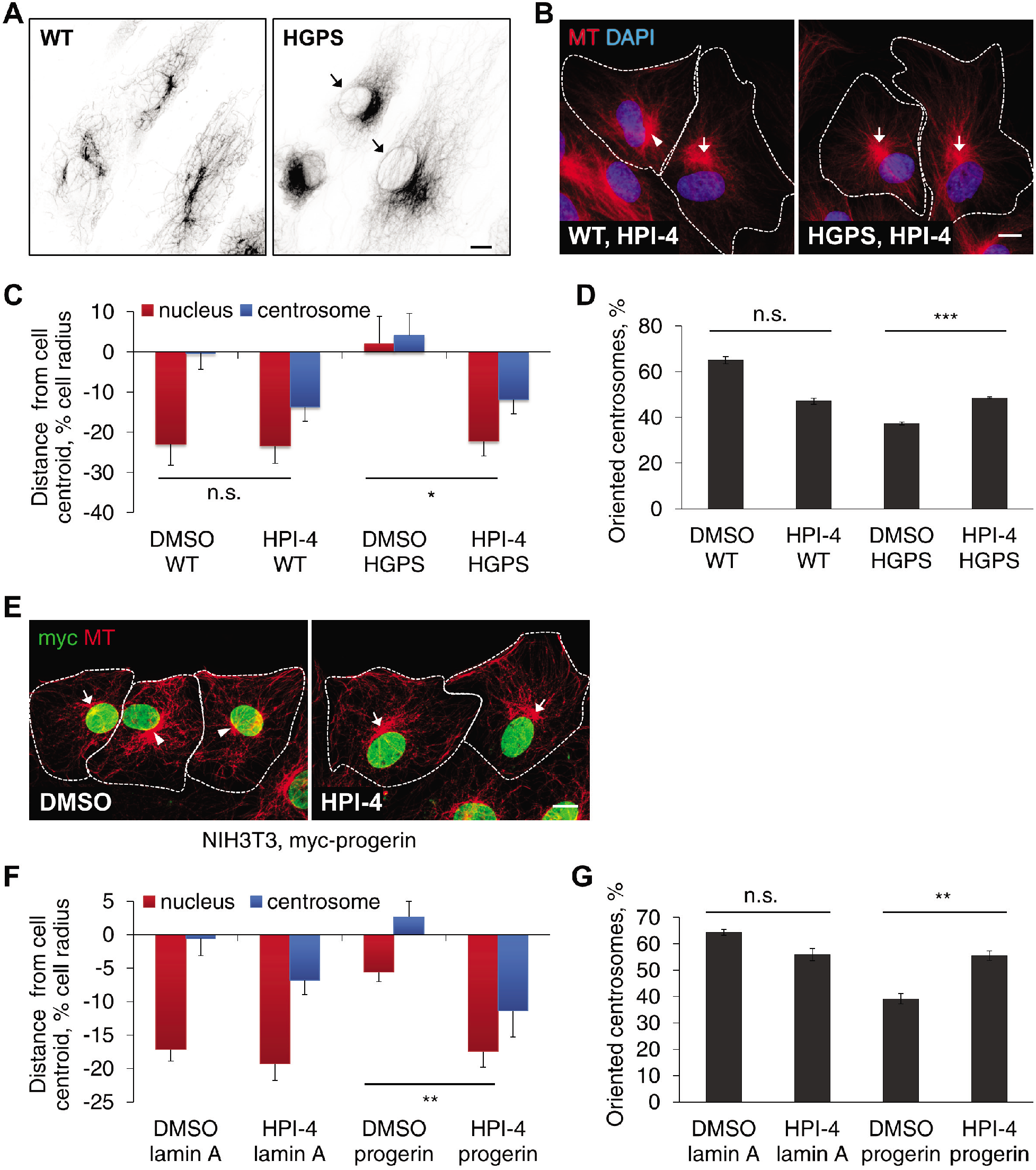
Cell polarity defects in HGPS fibroblasts are mediated by the microtubule cytoskeleton. *(A)* Representative images of microtubules (MT) and nuclei (DAPI) in WT (left) and HGPS (right) fibroblasts treated with 10 μM nocodazole for 2 h. Inverted contrast is shown. Nocodazole resistant microtubules surrounding nuclei (arrows) were observed more frequently in HGPS fibroblasts (29.5%, *n* = 670 cells, *N* = 2) compared to in WT fibroblasts (0.20%, *n* = 1000 cells, *N* = 2). *(B)* Representative images of microtubules (MT) and nuclei (DAPI) in WT (left) and HGPS (right) fibroblasts pretreated with 5 μM HPI-4 for 1 h and stimulated with LPA. Arrows, oriented centrosomes; arrowheads, unoriented centrosomes; dotted lines, cell borders. *(C,D)* Quantification of nuclear and centrosomal positions *(C)* and centrosome orientation *(D)* in LPA-stimulated WT and HGPS fibroblasts pretreated with vehicle (DMSO) or 5 μM HPI-4 for 1 h. *(E)* Images of microtubules (MT) and myc-progerin in NIH3T3 fibroblasts stably expressing myc-progerin and treated with DMSO or 10 μM HPI-4 for 1 h before LpA stimulation. Arrows, arrowheads and dotted outline are as in B. *(F,G)* Quantification of nuclear and centrosome positions *(F)* and centrosome orientation *(G)* in LPA-stimulated NIH3T3 fibroblasts expressing myc-tagged lamin A or progerin and pretreated with either DMSO or 10 μM HPI-4 for 1 h. Data are mean ± SEM from *n* ≥ 3 experiments (n > 90 cells). n.s. *p* > 0.05, \**p* < 0.05, \*\**p* < 0.01, \*\*\**p* < 0.001 by t-test. Bars, A,B,E: 10 μm.

### Fibroblasts from aged individuals exhibit similar nuclear movement and cell polarity defects as HGPS fibroblasts

As in HGPS (7, 28), nuclear architecture and cell migration are altered by physiological aging (5, 31, 32), but whether cell polarity is affected in aged migratory cells is unknown. We therefore examined centrosome orientation and nuclear movement in fibroblasts of comparable early passages from young (3-10 years old) and aged (87-90 years old) human males. Rearward nuclear movement and centrosome orientation stimulated by LPA occurred in fibroblasts from young but not aged individuals (Fig. 6A-C). To determine the age at which centrosome orientation becomes defective, we examined fibroblasts from a broader range of ages (SI Appendix, Table S4, S5). Centrosome orientation was normal in fibroblasts from both male and female subjects up to 60 years of age but was impaired in fibroblasts from virtually all those older than 60 years (Fig. 6D).

**Figure 6.**
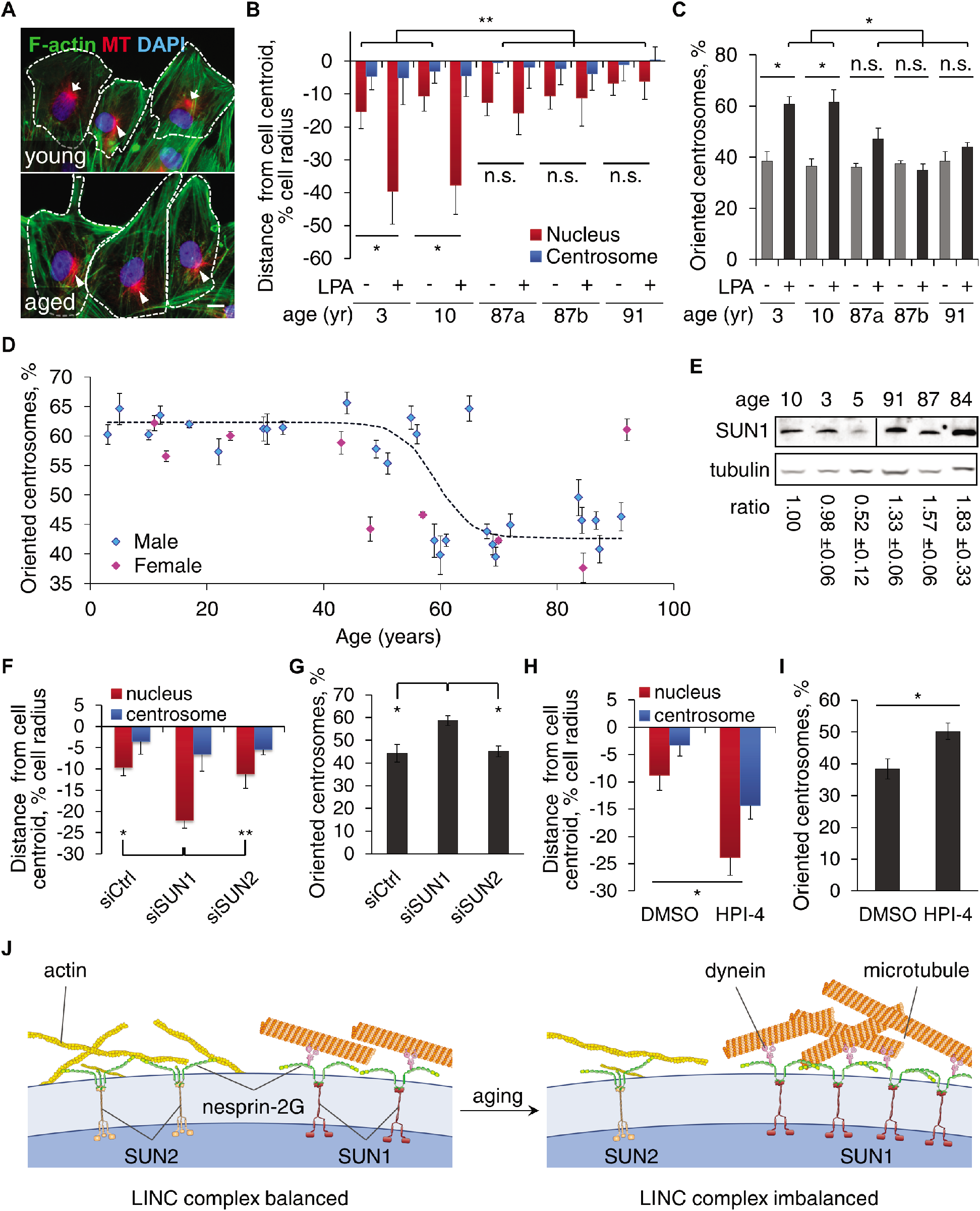
Fibroblasts from aged individuals exhibit cell polarity defects that are ameliorated by SUN1 knockdown and dynein inhibitor. *(A)* Representative images of F-actin, microtubules (MT) and nuclei (DAPI) in LPA-stimulated fibroblasts from young (10 yr) and aged (87 yr) male individuals. Arrows, oriented centrosomes; arrowheads, unoriented centrosomes; dotted outlines, cell borders. Bar, 10 μm. *(B,C)* Quantification of nuclear and centrosomal positions *(B)* and centrosome orientation *(C)* in LPA-stimulated fibroblasts from young and aged male individuals (87a and 87b are different individuals). *(D)* Quantification of centrosome orientation after LPA stimulation of fibroblasts from males (blue) and females (pink) of different ages. The trend line is a logistic regression curve fitted to the results for male fibroblasts (*p* < 0.0001, Pearson’s χ^2^ test). *(E)* Immunoblots of SUN1 and tubulin in fibroblasts from young and aged individuals. The SUN1/tubulin ratio is mean +/- SEM from 3 experiments and is normalized to the leftmost sample (*p* < 0.05 by t-test, young versus aged cells). *(F,G)* Quantification of nuclear and centrosomal positions *(F)* and centrosome orientation *(G)* in LPA-stimulated fibroblasts (87 yr) treated with indicated siRNAs. *(H,I)* Quantification of nuclear and centrosome positions *(H)* and centrosome orientation *(I)* in LPA-stimulated aged (87 yr) fibroblasts treated with vehicle (DMSO) or 5 μM HPI-4 for 1 h. Data are mean ± SEM from *n* ≥ 3 experiments (n > 90 cells). n.s. *p* > 0.05, \**p* < 0.05, \*\**p* < 0.01 by t-test. *(J)* Model for the imbalanced engagement of the nucleus with the cytoskeleton in fibroblasts from HGPS and aged individuals.

The inability of cells from aged individuals to polarize did not reflect a general defect in LPA-mediated signaling, as LPA-stimulated the assembly of actin stress fibers in these cells (see Fig. 6A), one of the canonical responses to LPA (33). These polarity defects did not correlate with dysmorphic nuclei (found in 6.9% of young and 8.7% of aged cells; ≥ 600 cells; *p* > 0.05 by χ^2^-test), proliferation (SI Appendix, Fig. S4A) or cellular senescence (SI Appendix, Fig. S4B, C), factors associated with aged cells as they are cultured for prolonged times (34). They also did not correlate with age- and passage-associated accumulation of DNA damage measured by phosphorylated H2AX (SI Appendix, Fig. S4D) (35). Additionally, although fibroblasts from young individuals accumulated markers of cellular senescence at later passages (SI Appendix, Fig. S4C), centrosome orientation in these cells was normal (SI Appendix, Fig. S4E). These results indicate that defects in nuclear movement and polarization are cell intrinsic phenotypes common to accelerated and physiological aging and are independent of other markers of cellular aging.

### Cell polarity defects in fibroblasts from aged individuals are rescued by reducing microtubule coupling to the nucleus

We detected progerin expression at low levels in fibroblasts from both young and aged individuals, but similar to a previous report, we did not detect a consistent difference between the samples (SI Appendix, Fig. S5). We next checked whether SUN1 was upregulated in fibroblasts from aged individuals as in HGPS fibroblasts. In whole cell lysates of fibroblasts from aged individuals, SUN1 expression was significantly elevated compared to lysates from fibroblasts from young individuals (Fig. 6E). These results suggested that the polarity defects in aged cells may have the same mechanistic basis as in the HGPS fibroblasts. Indeed, knock down of SUN1 (but not SUN2) rescued the polarity defects in aged fibroblasts (Fig. 6F, G). Similarly, HPI-4 treatment restored nuclear movement, and partially rescued centrosome orientation (Fig. 6H, I). These results indicate that elevated SUN1 expression is responsible for the cell polarity defects in fibroblasts from children with HGPS and physiologically aged individuals.

## Discussion

Establishing and maintaining polarity are essential for many cellular functions, including productive migration, which is critical for tissue renewal and repair (36). Reduced cell migration is an aging phenotype and has been proposed to contribute to some of the symptoms of HGPS (7, 28, 31, 32). Here we show that in fibroblasts from HGPS and aged individuals nuclear engagement with microtubules is favored compared to actin filaments, leading to an imbalance in the two systems and defects in cell polarization (Fig. 6J).

Our data suggest that the primary effect of progerin on actin-dependent nuclear movement and cell polarization is mediated by increased SUN1 levels. Elevated progerin expression increases SUN1 accumulation, most likely because of SUN1’s propensity to interact strongly with it (20). Elevated SUN1 is likely to have two consequences on actin-dependent nuclear movement. First, the preference of SUN1-nesprin-2G LINC complexes for microtubules (22) would make it more difficult for the nucleus to be moved by actin filaments. This would produce slipping TAN lines not because of an inability for them to be anchored as in lamin A null cells (19), but because the nucleus itself is excessively anchored. Consistent with this conclusion, we observed normal actin-dependent nuclear movement by both reducing SUN1 levels and by inhibiting dynein. Elevated SUN1 may also actively compete with SUN2 for a limited pool of nesprin-2G, thus limiting the formation of SUN2-nesprin-2G complexes. This idea is supported by the reduced nesprin-2-mediated actomyosin tension on the nuclear envelope measured in HGPS fibroblasts (37).

Although SUN1 is upregulated in fibroblasts from aged individuals, this does not seem to be due to upregulated progerin expression. We did not find a strong correlation between age and progerin expression in fibroblasts from normal individuals, consistent with a previous report (5). This suggests that either the aged cells are more sensitive to the deleterious effects of progerin (5) or that other age-related factors may enhance SUN1 accumulation.

We identify microtubules as a contributing factor to the cell polarity defects induced by progerin. Excessive microtubule coupling to the nucleus prevents its movement by actin and may also cause the actin flow phenotypes. The role of the microtubule cytoskeleton in HGPS is largely unexplored, although inhibition of the histone acetylase NAT10 appears to ameliorate nuclear defects in HGPS fibroblasts by reorganizing microtubules (38). Combined with our results, this suggests that targeting the microtubule cytoskeleton may be a possible treatment for HGPS.

It has been an ongoing debate whether the study of HGPS is relevant to our understanding of physiological aging (39, 40). Our findings strongly support the idea that at least some cellular and molecular defects in HGPS are shared with physiological aging. We observed both similar cellular defects and underlying molecular alterations in HGPS fibroblasts and fibroblasts from aged individuals. As other markers of aging do not correlate with the cell polarity defect, it is possible that the imbalance in the LINC complexes and/or the cell polarity defects precedes some of them. For example, the LINC complex has been implicated in DNA repair (41) and defects in DNA repair and other chromatin markers are dependent on progerin expression in HGPS and normal aged cells (5). Future research may determine whether there is a relationship between LINC complex imbalance, cell polarization defects and chromatin alterations in aging.

## Materials and methods

Primary fibroblasts were from Coriell Cell Repositories (Camden, NJ) and The Progeria Research Foundation (Peabody, MA). The nuclear movement assay, actin flow analysis and FRAP analysis were performed as previously described (18, 26). Other materials and experimental details are described in SI Materials and Methods.

## Supporting information

## Acknowledgments

Research reported in this publication was supported by NIH grants GM099481 to G.G.G. and AR068636 to G.G.G. and H.J.W. The authors declare no competing interests.

## Author Contributions

G.G.G. and H.J.W. conceived of the project and helped to analyze and interpret the data. W.C. designed and performed most of the experiments, analyzed data and prepared figures. Y.W. designed, performed and analyzed all FRAP experiments and experiments in SI Appendix, Fig. S2A,B. Y.W. and C.Ö. also contributed to data in Fig. 2A-C. G.W.G.L. did initial experiments on nuclear movement in progerin expressing NIH3T3 fibroblasts. C.Ö. made some of the cDNA constructs. W.C., G.G.G., and H.J.W. wrote the paper. All authors edited the manuscript.

