## Supplementary material for "Imbalanced Nucleocytoskeletal Connections Create Common Polarity Defects in Progeria and Physiological Aging"

**Supplemental material and methods**

**Reagents**

LPA was from Avanti Polar Lipids (Alabaster, AL). Dye-conjugated phalloidin was from Thermo Fisher Scientific (Waltham, MA). Unless noted, all other chemicals were from Sigma-Aldrich (St. Louis, MO). Lifeact-mCherry, EGFP-mN2G, EGFP-SUN1, EGFP-SUN2, EGFP-lamin A, EGFP-progerin, EGFP-progerin-SSIM, EGFP-emerin and EGFP-LAP1 were previously described (1-4). Dr. Arnoud Sonnenberg (Netherlands Cancer Institute, Amsterdam, Netherlands) provided EGFP-tagged nesprin-3α and nesprin-3β cDNA constructs and Dr. Kyle J. Roux (Sanford Children's Health Research Center, Sioux Falls, SD) provided a EGFP-nesprin-4 cDNA construct. EGFP-lamin B1 cDNA was cloned into pEEGFP-C1p, RFP-tagged lamin A and progerin cDNAs into RFP-C1 and Myc-tagged SUN1 and SUN2 cDNA into pMSCV vector. Pericentrin mouse and rabbit antibodies were from BD Transduction Laboratories (San Jose, CA) and Covance (Princeton, NJ), respectively. Tyrosinated α-tubulin rat monoclonal antibody (YL1/2) was from the European Collection of Animal Cell Cultures (Salisbury, UK). β-actin (clone C4) mouse antibody, c-myc rabbit antibody and lamin A/C rabbit antibody were from Santa Cruz Biotechnology (Santa Cruz, CA). Lamin A/C mouse antibody (MANLAC1) was from Dr. Glenn Morris (MDA Monoclonal Antibody Resource at the Wolfson Centre for Inherited Neuromuscular Disease, Oswestry, UK). SUN1 rabbit antibody was provided by Dr. Sue Shackleton (University of Leicester, UK), as well as from Sigma-Aldrich and Abcam (Cambridge, MA). Dr. Brian Burke (Institute of Medical Biology, Singapore) provided SUN1 mouse antibody. SUN2 and γH2AX rabbit antibodies were from Abcam. HDJ-2 mouse antibody was from Lab Vision (Fremont, CA). GAPDH mouse antibody was from Ambion (Austin, TX). Progerin mouse antibody was from Abcam. GFP chicken antibody was from Millipore Sigma (Burlington, MA). Myc mouse antibody was from Sigma-Aldrich. All secondary antibodies were from Jackson ImmunoResearch (West Grove, PA). Ultra-sensitive enhanced chemiluminescent HRP substrate was from Thermo Fisher Scientific.

**Cell culture and drug treatment**

Serum-free medium (SFM) is Dulbecco’s modified Eagle medium (Corning, Oneonta, NY) containing 10 mM HEPES (pH 7.4), penicillin and streptomycin (Thermo Fisher Scientific). NIH3T3 fibroblasts were from ATCC (Manassas, VA) and were cultured in SFM with 10% calf serum (Gemini, West Sacramento, CA). HEK293T cells (ATCC) and all human fibroblasts (Coriell Cell Repositories, Camden, NJ or The Progeria Research Foundation, Peabody, MA) were cultured in SFM with 15% fetal bovine serum (Gemini). NIH3T3 and human fibroblasts were only used for 6-8 passages except when the effect of passage on nuclear movement and senescence was tested. To serum starve cells, confluent monolayers were washed three times with SFM and maintained in SFM for 2 (NIH3T3) or 3 d (human fibroblasts). Serum-starved cells were wounded and nuclear movement was stimulated with 10 μM LPA for 45 min to detect TAN lines or 2 h to measure nuclear movement and centrosome orientation. FTI-277, when added, was used at 2.5 μM and the medium was changed daily. FTI-227 treatments were 48 h for nuclear movement assays and 72 h for FRAP assays.

**Transfection and infection**

For live cell imaging, plasmid DNA (10 μg/ml) diluted in 140 mM KCl, 10 mM HEPES (pH 7.4) was pressure microinjected into nuclei of cells at the wounded edge of monolayer cultures and allowed to express for 1 h before adding 20 μM LPA to stimulate nuclear movement. siRNAs (siScramble: UUCUCCGAACGUGUCACGU; siSUN1-1: CAAUCAGUGCGGUUGGUGA; siSUN1-1: GAAACUUACGAAACCAAAA; siSUN2: GGAAAUCCAGCAACAUGAA) were from

Shanghai GenePharma Co., Ltd (Shanghai, China) and were transfected at 20 µM using Lipofectamine RNAiMAX (Thermo Fisher Scientific). Stable cell lines were generated by infecting cells with retrovirus, produced in HEK293T cells, in the presence of 2 µg/ml polybrene (Millipore, Billerica, MA).

**FRAP**

Fibroblasts were cultured on chambered cover glasses and transfected with GFP-tagged constructs using Lipofectamine PLUS (Thermo Fisher Scientific). Experiments were performed on a Zeiss LSM 510 META confocal laser scanning system (Carl Zeiss, Oberkochen, Germany) attached to a Zeiss Axiovert 200 microscope with spectral resolution of fluorescence labels using the 488 nm line with a 30mW argon laser in conjunction with a Plan-Neofluar 40x/1.3 oil objective. A selected area with same size at each nucleus was photobleached at 50% laser power (100% transmission) for 2 iterations and the fluorescence recovery was monitored by scanning at low power (2.1% transmission) in 3 s intervals for the first 30 micrographs and 5 or 8 s intervals for another 70 micrographs. We also performed FRAP using a Nikon A1R MP multiphoton confocal microscope with imaging software NIS-Elements and environment control at 37°C and 5% $CO_2$ (Nikon, Melville, NY). The average fluorescence intensity was measured in the region of interest and normalized to the change in total fluorescence as $I_{rel} = T_0 I_t / T_t I_0$ as described previously (2). The halftime of recovery was calculated as $t_{1/2} = ln2 \times \frac{-1}{slope}$ using data taken from the first 30 s after bleaching as reported previously (2).

**Microscopy**

For immunofluorescence microscopy, cells on coverslips were fixed in 4% paraformaldehyde (Electron Microscopy Sciences, Hatfield, PA), permeabilized in phosphate-buffered saline containing 5% normal goat serum and 0.3% Triton X-100 and stained with standard techniques. Coverslips were mounted with Fluoromount-G (Southern Biotech, Birmingham, AL). Fluorescence and phase contrast images were acquired with either a 40X PlanApo objective (NA 1.0) and a CoolSNAP HQ CCD camera on a Nikon TE300 inverted microscope controlled by Metamorph (Molecular Devices, Sunnyvale, CA), or with a Zeiss LSM 510 META confocal laser scanning system attached to a Zeiss Axiovert 200 inverted microscope. Movies of live cell fluorescent proteins were acquired at 37˚C (5 min per frame) with a 60X PlanApo objective (NA 1.49) and an iXon X3 CCD camera on a Nikon Eclipse Ti microscope controlled by NIS-Elements. Microscopic images and movies were processed using Fiji.

**Data analysis**

Nuclear and centrosomal positions were measured using CellPlot as previously described (5) and analyzed and plotted using Excel (Microsoft, Redmond, WA). The velocities of actin retrograde flow were calculated from the slopes of lines in kymographs generated using NIS-Elements and Fiji as previously described (6). We developed custom software (available upon request) to quantify background-subtracted immunofluorescence staining of nuclear proteins (in Fig. 5B). Quantification of immunoblot signals was done using Fiji with background subtraction. Statistical analysis was performed in Microsoft Excel. Unpaired two-tailed Student's t-tests were used to calculate $p$ values. The trend line in Fig. 6D was the logistic regression curve fitted to the results from male fibroblasts, with 0 assigned to centrosome orientation values < 50% and 1 for values ≥ 50%. The probabilities of failed centrosome orientation ($p$) were calculated from the fit curve and normalized to the range of centrosome orientation values using the equation: $min \times p + max \times (1 - p)$. Here $min$ is the average of 10 lowest values and $max$ is the average of 10 highest values.

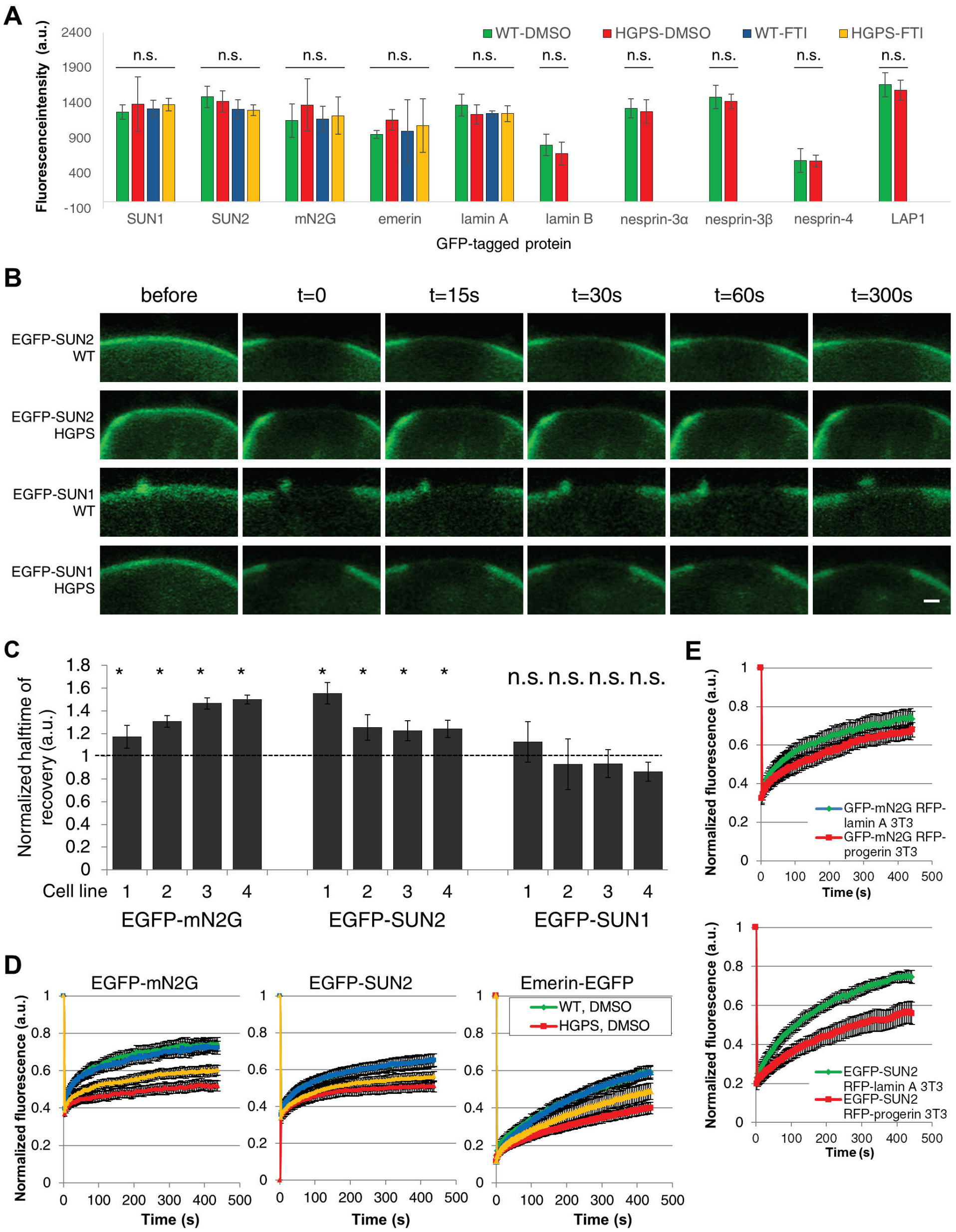


**Fig. S1. Progerin expression reduces the diffusional mobilities of a subset of nuclear envelope proteins.**

*(A)* Quantification of GFP intensities in cell used in FRAP experiments. n.s. $p > 0.05$ by one-way ANOVA test (from SUN1 to lamin A, 3 experiments, $n > 50$ cells) or by Student's t-test (from lamin B to LAP1, 3 experiments, $n > 30$ cells). *(B)* Representative images of nuclei from WT and HGPS fibroblasts expressing either EGFP-SUN2 or EGFP-SUN1 before and at the indicated times after photobleaching. Bar, 1 µm. *(C)* Normalized $t_{1/2}$ of FRAP of EGFP-tagged mN2G, SUN2, and SUN1 in four pairs of HGPS fibroblasts and sex- and age-matched controls. Statistical tests were versus controls. *(D)* Normalized fluorescence recovery after photobleaching for EGFP-tagged nesprin-2G, SUN2, and emerin, in nuclei of normal fibroblasts (WT: green, DMSO-treated; blue, FTI-treated) and HGPS fibroblasts (HGPS: red, DMSO-treated; orange, FTI-treated). Cells were treated with either DMSO or 2.5 µM FTI-277 for 72 h before FRAP. *(E)* Normalized fluorescence recovery after photobleaching for EGFP-mN2G and EGFP-SUN2 in nuclei of NIH3T3 fibroblasts expressing RFP-tagged lamin A (green) or progerin (red). Data are represented as the mean ± SEM from ≥ 3 experiments ($n > 15$ cells).

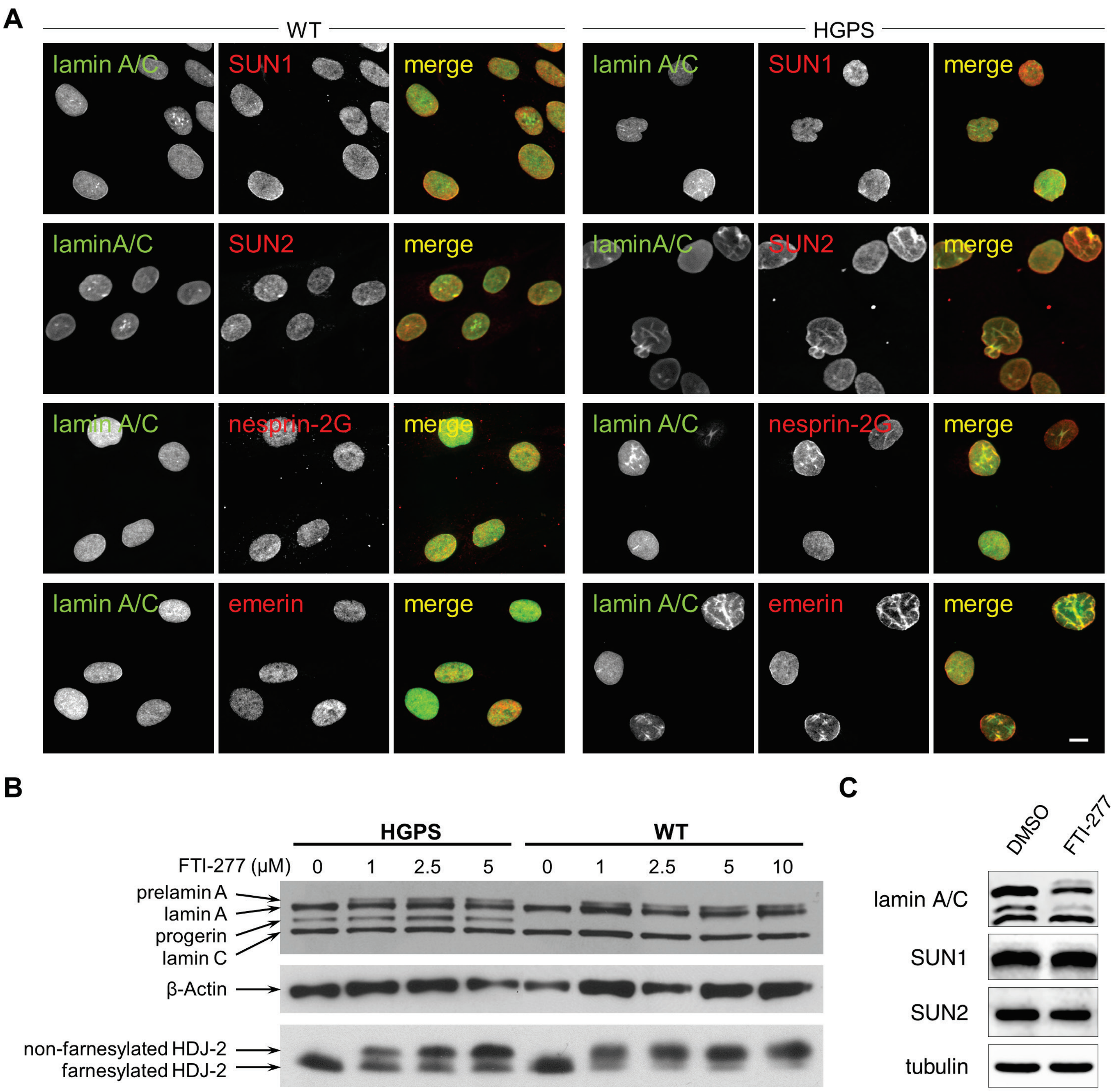


**Fig. S2. Localization of emerin, nesprin-2G, SUN1, and SUN2 are similar in control and HGPS fibroblasts and FTI-277 blocks protein farnesylation.**
*(A)* Micrographs showing immunofluorescence labeling of lamin A/C, SUN1, SUN2, nesprin-2G (N2G) and emerin in fibroblasts from a control individual (WT) and an individual with HGPS. Bar, 10 μm. *(B)* Immunoblot showing that increasing concentrations of FTI-277 interfered with prelamin A processing and caused accumulation of prelamin A in HGPS and control (WT) fibroblasts (top). β-actin (middle) is a loading control. HDJ-2 (bottom) is another farnesylation substrate and migrates slower when it is non-farnesylated. *(C)* Immunoblot showing that the levels of SUN proteins are not affected by 48 hours of FTI treatment.

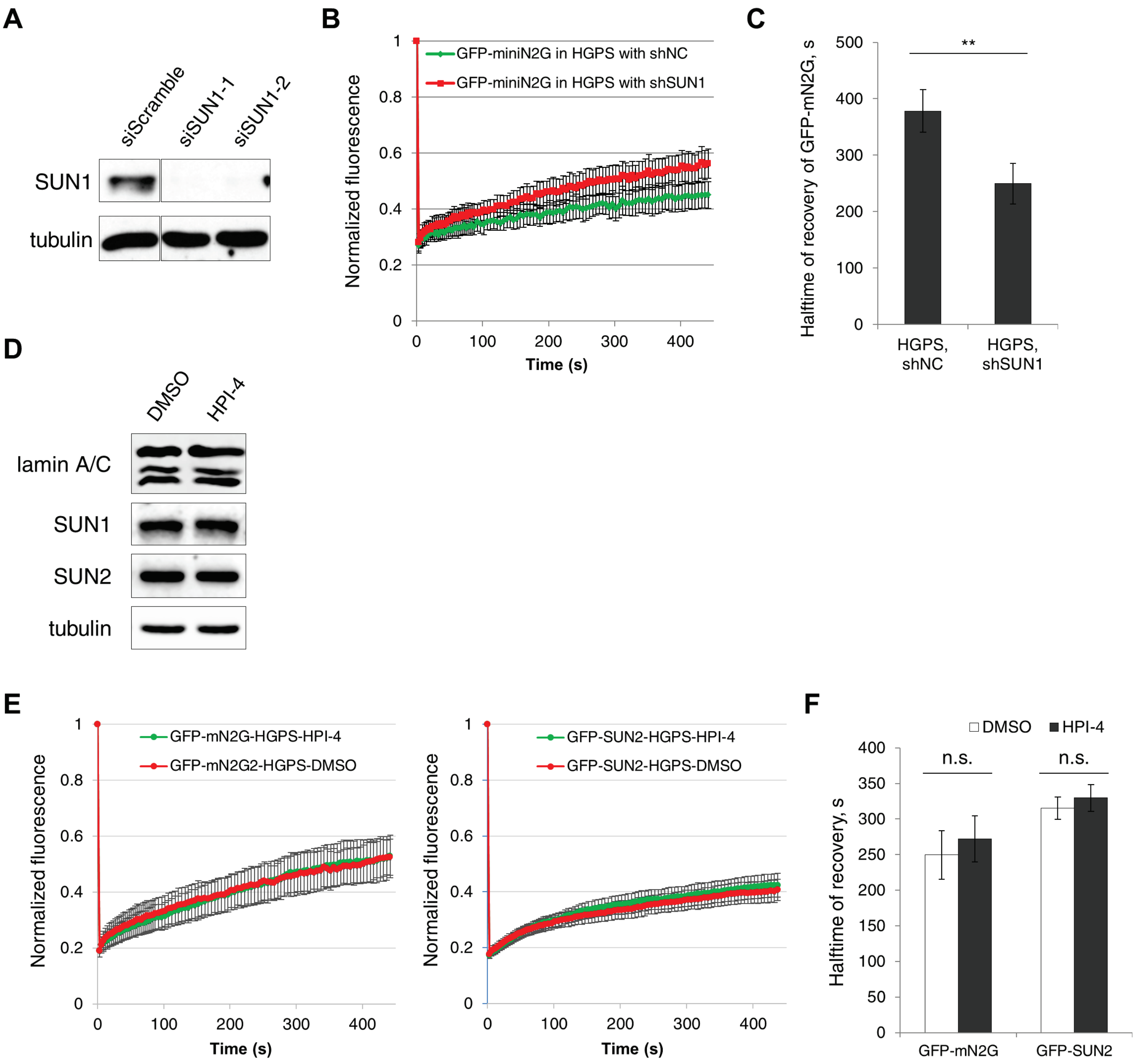


**Fig. S3. SUN1 Depletion increased diffusional mobility of GFP-mN2G in HGPS fibroblasts.** *(A)* Immunoblot showing that two distinct SUN1 siRNAs efficiently depleted SUN1 in human fibroblasts. Tubulin is a loading control. *(B)* Normalized FRAP for EGFP-mN2G in HGPS fibroblasts expressing control (shNC, green) and SUN1 specific (shSUN1, red) shRNAs ($N$ = 3 experiments, $n$ > 15 cells). *(C)* The $t_{1/2}$ of EGFP-mN2G for data in B. *(D)* Immunoblot showing that the levels of SUN proteins are not affected by 3 hours of 10 μM HPI-4 treatment. *(E)* Normalized FRAP for EGFP-mN2G and EGFP-SUN2 in HGPS fibroblasts treated with DMSO or 10 μM HPI-4. ($N$ = 3 experiments, $n$ > 15 cells). *(F)* The $t_{1/2}$ of EGFP-tagged proteins for data in E. n.s. p>0.05; ** $p$ < 0.01 by Student's t-test. Data are represented as the mean ± SEM.

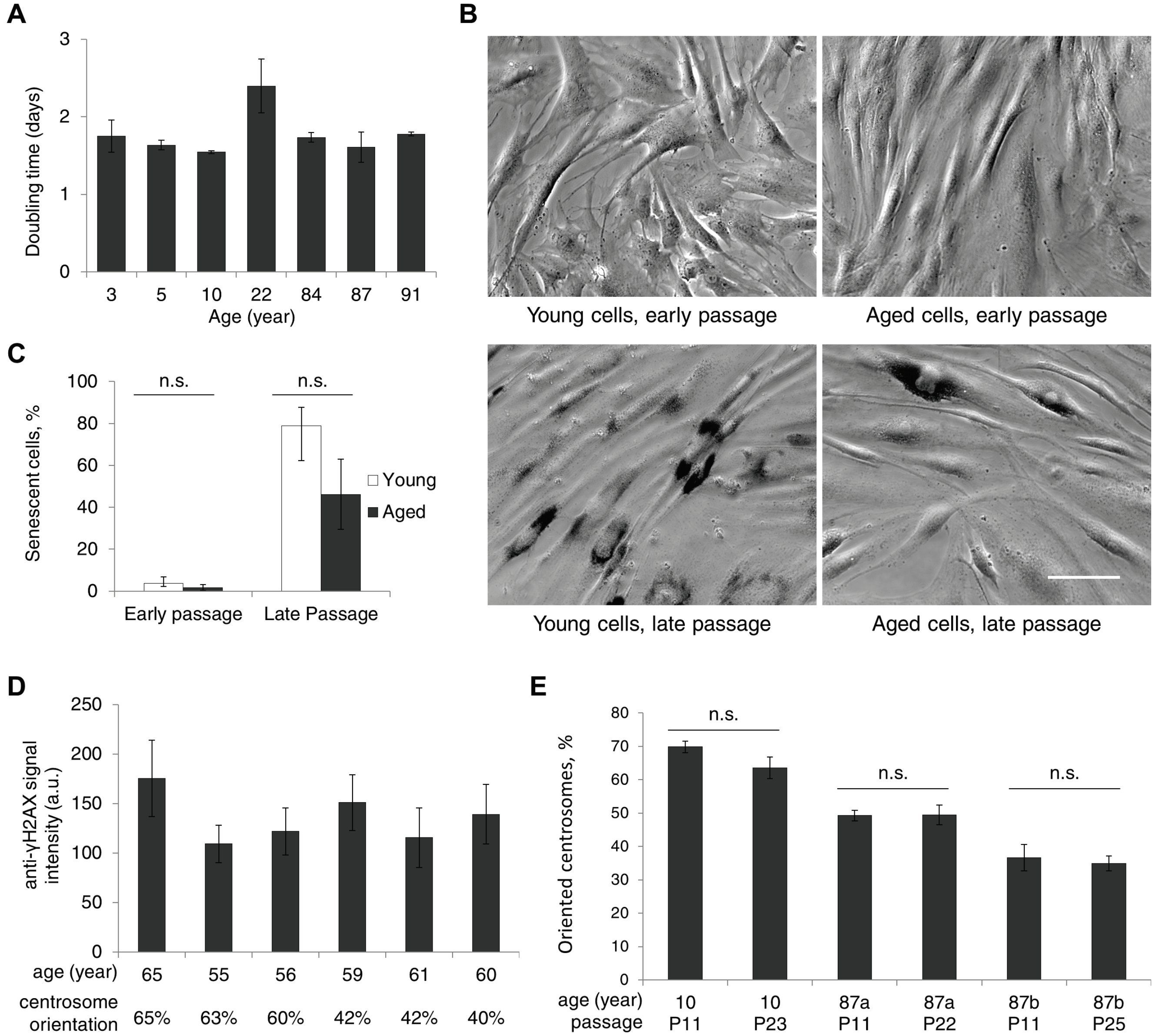


**Fig. S4. Cell polarity defects in fibroblasts from aged individuals do not correlate with proliferation, senescence or DNA damage.**

*(A)* Growth rates of fibroblasts from individuals vary without clear correlation with their ages of sampling. The fibroblasts from young and aged individuals used in nuclear movement assays exhibited similar growth rates, as measured by doubling times. There was no significant difference in growth rates between young (3-10) and old (84-91) fibroblasts by t-test ($N = 2$ experiments). *(B)* Representative images of early (≤ 15) and late (> 20) passages of fibroblasts from both young and aged individuals stained for senescence β-galactosidase activity (black staining). Bar, 50 μm. *(C)* Quantification of cellular senescence for the cells shown in B. ($N = 3$ experiments, $n > 150$ cells). *(D)* Quantification of DNA double strand breaks as measured by γH2AX signal in fibroblasts from six individuals with similar ages at sampling ($n > 150$ cells). The level of centrosome orientation is shown below. Student's t-test showed that there was no significant difference between fibroblasts with normal (left three) compared to defective (right three) centrosome orientation ($p > 0.99$). *(E)* Centrosome orientation was similar at early and late passage in fibroblasts from either young or aged individuals. ($N = 2$ experiments, $n > 150$ cells). Data are represented as the mean ± SEM. n.s. $p > 0.05$ by Student's t-test.

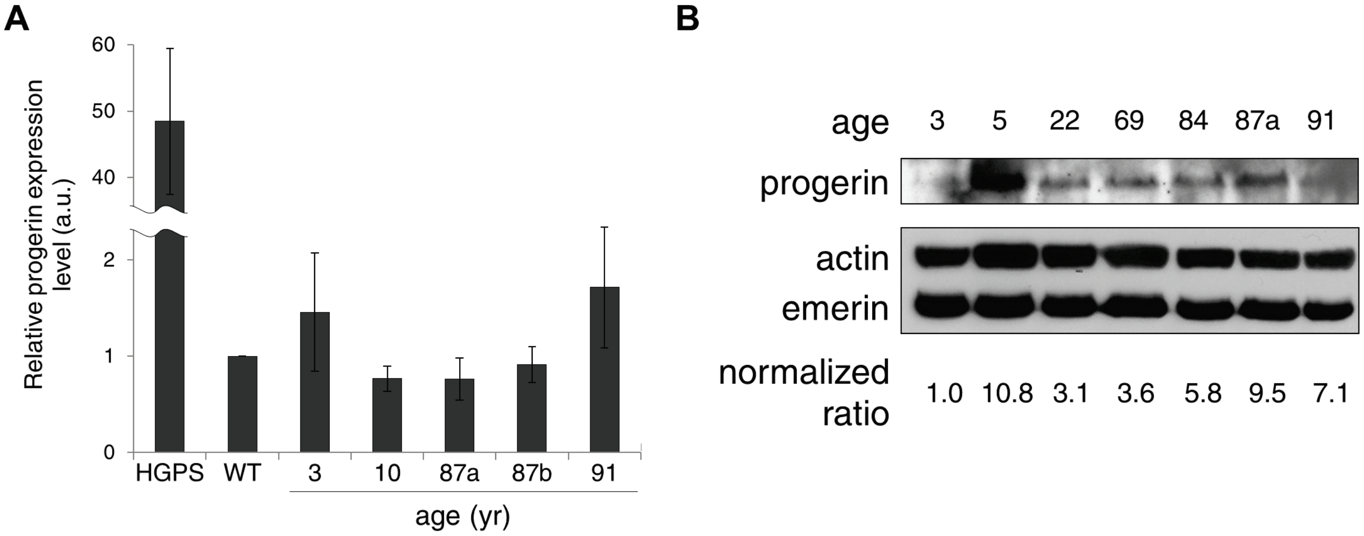


**Fig. S5. Progerin level in fibroblast do not correlate with age.**
(A) Quantification of progerin mRNA in dermal fibroblasts from young and aged individuals by RT-PCR. Fibroblasts from a subject with HGPS served as a positive control. There is no significant difference in progerin mRNA levels in fibroblasts from young and aged individuals by t-test. Error bars, SEM from n = 3 measurements. (B) Progerin protein (top) was detected in whole cell lysates (see Methods) of fibroblasts from both young and aged individuals. Actin and emerin (bottom) served as loading control. The ratio is progerin/actin normalized to first sample.

### Supplemental tables

**Table S1. Pairs of dermal fibroblast lines from control individuals (WT) and children with HGPS.**

| Name | ID | Sex | Age (yr) | Source | Note |
|---|---|---|---|---|---|
| **WT1** | GM00316 | male | 12 | CCR* | |
| **HGPS1** | AG11498 | male | 14 | CCR | |
| **WT2** | GM00498 | male | 3 | CCR | |
| **HGPS2** | AG06917 | male | 3 | CCR | |
| **WT3** | GM01652 | female | 11 | CCR | |
| **HGPS3** | AG01972 | female | 14 | CCR | |
| **WT4** | GM00038 | female | 9 | CCR | |
| **HGPS4** | AG11513 | female | 8 | CCR | |
| **WT5** | HGFDFN 168 | male | 40 | PRF† | father of HGADFN 167 |
| **HGPS5** | HGADFN 167 | male | 8 | PRF | son of HGFDFN 168 |
| **WT6** | HGMDFN 371 | female | 45 | PRF | mother of HGMDFN 370 |
| **HGPS6** | HGADFN 370 | female | 10 | PRF | daughter of HGMDFN 371 |

* CCR: Coriell Cell Repositories
† PRF: Progeria Research Foundation

**Table S2. Summary of $t_{1/2}$ in fibroblasts derived from matched controls (WT) and subjects with HGPS for FRAP experiments (Figure 1B).**

| Cell lines | $t_{1/2}$ (s)* EGFP-SUN2 | $t_{1/2}$ (s) EGFP-SUN1 | $t_{1/2}$ (s) EGFP-mN2G |
|---|---|---|---|
| WT1 | 112.7 ± 13.5 | 178.1 ± 25.2 | 79.2 ± 9.3 |
| HGPS1 | 175.3 ± 10.5 | 200.7 ± 31.6 | 92.9 ± 7.8 |
| WT2 | 138.2 ± 18.5 | 174.6 ± 17.7 | 88.2 ± 5.8 |
| HGPS2 | 173.3 ± 15.7 | 162.2 ± 39.1 | 115.4 ± 7.4 |
| WT3 | 140.5 ± 12.5 | 165.1 ± 28.9 | 103.3 ± 7.5 |
| HGPS3 | 172.2 ± 12.3 | 154.3 ± 20.4 | 151.5 ± 11.4 |
| WT4 | 145.0 ± 12.5 | 164.3 ± 22.6 | 102.0 ± 5.9 |
| HGPS4 | 180.0 ± 11.3 | 142.1 ± 13.8 | 153.0 ± 14.2 |

* $t_{1/2}$ is expressed as means ± standard errors from ≥ 3 experiments ($n$ > 15 cells).

**Table S3. Summary of $t_{1/2}$ in FRAP experiments (Figures 1C, 1D, S1, and S6).**

| EGFP-tagged protein | Treatment | $t_{1/2}$ (s)* | |
|---|---|---|---|
| | | WT | HGPS |
| SUN2 | DMSO | 147.2 ± 9.8 | 196.8 ± 10.9 |
| SUN2 | FTI-277 | 134.8 ± 6.2 | 164.3 ± 8.3 |
| mN2G | DMSO | 86.3 ± 10.2 | 184.1 ± 12.8 |
| mN2G | FTI-277 | 94.5 ± 10.0 | 142.2 ± 11.0 |
| Emerin | DMSO | 185.5 ± 11.6 | 269.0 ± 10.0 |
| Emerin | FTI-277 | 169.5 ± 11.6 | 210.1 ± 7.6 |
| **EGFP-tagged protein** | | **NIH3T3 expressing RFP-lamin A** | **NIH3T3 expressing RFP-progerin** |
| SUN2 | | 159.1 ± 11.1 | 260.6 ± 19.7 |
| mN2G | | 94.5 ± 12.2 | 148.6 ± 18.5 |
| **EGFP-tagged protein** | | **HGPS expressing shNC** | **HGPS expressing shSUN1** |
| mN2G | | 378.2 ± 37.6 | 249.7 ± 35.9 |

* $t_{1/2}$ is expressed as means ± standard errors from ≥ 3 experiments (*n* > 15 cells).

**Table S4. Dermal fibroblasts from apparently normal men used in this study, related to Figure 4.**

| Strain | Catalog ID[a] | Biopsy source | Race/Ethnicity | Age at biopsy (yr) | Population doubling level at freeze | Passage frozen |
|---|---|---|---|---|---|---|
| **3** | GM05565 | inguinal | Hispanic/Latino | 3 | | 5 |
| **5** | GM05381 | umbilical area | Black | 5 | | 3 |
| **10** | GM03348 | unspecified | Caucasian | 10 | | 7 |
| **12** | GM00316 | arm | Caucasian | 12 | | 12 |
| **17** | AG06234 | arm | Caucasian | 17 | 6.5 | 3 |
| **22** | AG11747 | arm | Caucasian | 22 | 16 | |
| **30a** | AG11242 | arm | Caucasian | 30 | 9 | |
| **30b** | AG13153 | arm | Caucasian | 30 | 5.54 | 6 |
| **33** | AG04438 | arm | Caucasian | 33 | 13 | |
| **44** | AG13156 | arm | Caucasian | 44 | 7 | 3 |
| **49** | AG11160 | arm | Caucasian | 49 | 6 | |
| **51** | AG07136 | arm | Caucasian | 51 | 9 | |
| **55** | AG12657 | arm | Caucasian | 55 | 5 | 2 |
| **56** | AG13292 | arm | Caucasian | 56 | 5 | 2 |
| **59** | AG06239 | arm | Caucasian | 59 | 8 | |
| **60** | AG05419 | arm | Caucasian | 60 | 19 | |
| **61** | AG11011 | arm | Caucasian | 61 | 7 | |
| **65** | AG04659 | arm | Caucasian | 65 | 9 | |
| **68** | AG16030 | arm | Caucasian | 68 | 6 | 3 |
| **69a** | AG05095 | arm | Caucasian | 69 | 8 | |
| **69b** | AG05099 | arm | Caucasian | 69 | 7 | |
| **72** | AG04455 | arm | Caucasian | 72 | 9 | |
| **84a** | AG05274 | arm | Caucasian | 84 | 14 | |
| **84b** | AG11730 | arm | Caucasian | 84 | 5.97 | 4 |
| **87a** | AG10884 | arm | Caucasian | 87 | 7 | |
| **87b** | AG05248 | arm | Caucasian | 87 | 16 | |
| **91** | AG07725 | arm | Caucasian | 91 | 11 | |

[a]Source: Coriell Cell Repositories.

**Table S5. Dermal fibroblasts from apparently normal women used in this study, related to Figure 4D.**

| Catalog ID[a] | Biopsy source | Race/Ethnicity | Age at biopsy (yr) | Population doubling level at freeze | Passage frozen |
|---|---|---|---|---|---|
| **GM02036** | unspecified | Caucasian | 11 | 6.21 | 13 |
| **GM01651** | arm | Caucasian | 13 | 7.09 | 11 |
| **AG11732** | arm | Caucasian | 24 | 7.11 | |
| **AG11745** | arm | Caucasian | 43 | 8 | |
| **AG11793** | arm | Caucasian | 48 | 8 | |
| **AG11564** | arm | Caucasian | 57 | 9 | |
| **AG11733** | arm | Caucasian | 70 | 8 | |
| **AG11725** | arm | Caucasian | 84 | 8 | |
| **AG09602** | arm | Caucasian | 92 | 5.48 | 2 |

[a]Source: Coriell Cell Repositories.

### Description of Supplemental Videos

#### Video S1

Time-lapse movie of LPA-stimulated actin cable movement in a mCherry-LifeAct and EGFP-progerin expressing NIH3T3 cell (panels presented in Fig. 3C). Wound edge is at the top. Actin cables move rearward but the nucleus is immobile. Progerin accumulates on actin cables and appears to move together with them, suggesting that the lamina with progerin cannot resist the force from actin cables and fails to transduce force to move the nucleus. Time: h:min.

#### Video S2

Time-lapse movie of LPA-stimulated NIH3T3 fibroblast expressing mCherry-LifeAct and EGFP-lamin A (panels presented in Fig. 3E). Wound edge is at the top. Actin cables move rearward in an oriented fashion and their velocity is much faster than that in an EGFP-progerin expressing cell (see Video S3). Time: h:min.

#### Video S3

Time-lapse movie of LPA-stimulated NIH3T3 cell expressing mCherry-LifeAct and EGFP-progerin (panels presented in Fig. 3F). Wound edge is at the top. Actin cables move haphazardly and their velocity is slower compared to that in NIH3T3 fibroblasts expressing EGFP-lamin A (see Video S2) or EGFP-progerin-SSIM (see Fig. 3G), indicating that progerin expression also impacts actin flow. Time: h:min.
